# Long-Term Surveillance Reveals Establishment of *Aedes albopictus* in Eastern Nebraska, USA

**DOI:** 10.64898/2026.09.23.750200

**Authors:** Gwenyth Coleman, Jeff Hamik, Halie Smith, Tyler Mucha, Leslie C. Rault, Troy D. Anderson

## Abstract

*Aedes albopictus* (Skuse), the Asian tiger mosquito, is a highly competent arboviral vector whose range has expanded substantially across the United States over the past four decades. Despite predictive models placing Nebraska within the species’ climatically suitable range, its establishment status in the state has remained poorly characterized. Here, we report results from a nine-year mosquito surveillance program (2017–2025) conducted across 44 Nebraska counties in collaboration with the Nebraska Department of Health and Human Services. *Ae. albopictus* was detected in five counties, with sustained, annually increasing populations documented in Richardson, Douglas, and Lancaster counties. Richardson County recorded continuous detections during 2017– 2025, with proportional representation rising to 60.50% of collected mosquitoes by 2025. In Douglas and Lancaster counties, temporal advancement of first seasonal detection in 2024 and 2025 provide evidence consistent with successful overwintering rather than annual reintroduction. A cumulative degree-day model predicted adult emergence in mid-May across all county-year combinations, consistently preceding trap deployment by two to seven weeks and revealing a systematic early-season surveillance gap. Generalized linear mixed-effects models indicated that trap-level detection persistence, rather than urban location, was the primary predictor of yearly *Ae. albopictus* positivity, suggesting that current invasion dynamics are driven by focal source populations. These findings provide strong evidence for the establishment of *Ae. albopictus* in eastern Nebraska and highlight the need for earlier seasonal surveillance and standardized criteria to define establishment in northward-expanding vector populations.

## Introduction

*Aedes albopictus* (Skuse 1849), commonly known as the Asian tiger mosquito, is a highly competent vector of more than 20 arboviruses, including dengue, Zika, chikungunya, yellow fever, and West Nile viruses (Paupy et al. 2009, Torto and Tchouassi 2024). Unlike many mosquito species that restrict host-seeking activity to specific times of day, *Ae. albopictus* exhibits opportunistic feeding behavior throughout the day, making it an effective bridge vector between zoonotic arboviruses and human populations (Paupy et al. 2009, Fortuna et al. 2015). Of particular public health concern is its capacity for transovarial transmission of West Nile virus (WNV), whereby infected females pass the virus to their eggs, allowing the pathogen to persist through winter diapause and contributing to both long-term survival and long-distance dispersal under adverse environmental conditions (Zhang et al. 2022). In temperate regions, *Ae. albopictus* often serves as the primary vector for arboviral transmission (Gratz 2004). This is especially relevant in Nebraska, where WNV remains a recurring public health burden, with multiple human clinical cases reported annually (Nebraska Department of Health and Human Services 2026). While *Culex* mosquitoes are considered the primary enzootic vectors of WNV in the region, the potential establishment of *Ae. albopictus* presents an additional and understudied transmission risk.

The distribution of *Ae. albopictus* is strongly influenced by abiotic factors, particularly temperature. In the United States, populations are often limited by a -5°C isotherm during winter months (Nawrocki and Hawley 1987), and global distributions of the species broadly correlate with temperature gradients (Kraemer et al. 2015). A key mechanism enabling survival under unfavorable conditions is facultative diapause of *Ae. Albopictus* eggs (Lee et al. 2024), which allows populations to persist through periods of cold temperatures. Due to ongoing climate change, regions that were once unsuitable for certain mosquito species are becoming increasingly habitable (Wilke et al. 2021). Nebraska, for instance, has experienced a steady increase in average annual temperatures, particularly during winter months (Oglesby et al. 2015), potentially facilitating the overwintering and establishment of thermally sensitive species such as *Ae. albopictus*. In addition, the urban heat island effect and land cover changes associated with urbanization are known to increase mosquito presence in populated areas (Wiese et al. 2019, Ravasi et al. 2022), further creating conditions favorable for establishment.

Since its first detection in Texas in 1985, *Ae. albopictus* has expanded its range to more than 26 states, spreading both northward and westward (Hahn et al. 2017, Swan et al. 2022). Previous studies have detected *Ae. albopictus* across parts of the U.S. Midwest, though many of these detections represent rare or unsuccessful introduction events (Hall et al. 2022). However, with the confirmed establishment of *Ae. albopictus* in the neighboring states of Iowa and Missouri (Hahn et al. 2017, Claborn et al. 2018, Hall et al. 2022, Lee et al. 2026), and with predictive models placing Nebraska within the species’ projected range, there is an elevated likelihood that it could also establish permanent populations in the state.

Temperature-based predictive models offer a proactive approach to managing *Ae. albopictus* by identifying when adult emergence is likely to occur, enabling more targeted and timely control measures (Fonseca et al. 2013, Armstrong et al. 2017, Healy et al. 2019). Teng and Apperson (2000) developed the foundational degree-day model for *Ae. albopictus*, which was subsequently expanded by Healy et al. (2019) to incorporate thermal maximum thresholds that influence larval mortality. Despite availability of this modeling framework, its predictive utility has not been evaluated in the context of Nebraska’s climate, and the establishment status of *Ae. albopictus* in the state remains poorly characterized.

In this study, we (i) document spatial and temporal patterns of *Ae. albopictus* detection across Nebraska during 2017–2025, (ii) evaluate a cumulative degree-day model for predicting adult emergence under Nebraska climate conditions, and (iii) test whether urbanization and trap-level persistence predict detection probability.

## Materials and Methods

### Ae. albopictus surveillance

Surveillance data were obtained from the Nebraska Department of Health and Human Services (DHHS) Vector-Borne Disease Program. Mosquito collections were conducted annually during 2017 through 2025, with additional collaboration from the University of Nebraska-Lincoln in 2024 and 2025. Surveillance was conducted in 44 of Nebraska’s 93 counties, although sampling intensity varied by year due to personnel availability and logistical constraints.

Collections were conducted between Morbidity and Mortality Weekly Report (MMWR) weeks 22 and 39. Traps were deployed on a biweekly schedule, set after 16:00, and retrieved the following morning between 08:00 and 09:00.

Two trap types were used. The primary trap was a CO₂-baited CDC light trap, with dry ice serving as the CO₂ source. Beginning in 2024, the light component was removed to reduce non-target bycatch. When intensified surveillance was required, BG-Sentinel 2 traps (Biogents AG) were deployed for overnight collections, baited with both dry ice CO₂ and BG-Sweet Scent lure, which mimics lactic acid and other human skin volatiles.

### Mosquito sample processing and identification

Following collection, mosquito specimens were transported to DHHS’s laboratory for identification to species and sex. Specimens were identified under a stereomicroscope using *Mosquitoes of North America* as the primary taxonomic reference (Darsie and Ward 2005). Counts for each species were recorded per trap, along with associated metadata including trap deployment date, GPS coordinates (longitude and latitude), city, and county (Supp. Table S1).

### Cumulative degree-day model

Thermal development parameters for *Ae. albopictus* were derived from experimentally validated nonlinear development models described by Healy et al. (2019). The model incorporated a lower developmental threshold temperature (T_min_ = 10.5°C), an optimal developmental temperature (T_peak_= 32.35°C), and an upper developmental threshold (T_max_ = 36.2°C). The cumulative thermal requirement for development to adulthood was defined as 172.4 cumulative degree-days (CDD).

Daily degree-days (DD) were calculated using a nonlinear temperature-response function based on mean daily temperature (MDT) (Healy et al. 2019). Briefly, DD values were assigned using the equation [CDD = ∑ DD]. The conditional structure included: 1) if MDT < T_min_, then DD = 0, 2) if MDT > T_max_, then DD = 0, 3) if T_max_ > MDT > T_peak_, then DD = T_peak_, and 4) if T_peak_ > MDT > T_min_, then DD = MDT − T_min_ (Healy et al. 2019). Daily DD values were then summed within each calendar year to generate CDD estimates, which were used to predict the timing of adult emergence.

### Temperature Data Acquisition

Daily air temperature data were obtained from the National Aeronautics and Space Administration (NASA) Langley Research Center (LaRC) Prediction of Worldwide Energy Resources (POWER) system through the Application Programming Interface (API) (NASA LaRC 2026). Data were retrieved using geographic coordinates representative of each study region: Douglas County (41.31° N, −96.20° W), Lancaster County (40.8471° N, −96.6638° W), and Richardson County (40.0833° N, −95.7000° W).

Extracted variables included daily mean air temperature at 2 m height, as well as daily minimum and maximum air temperatures. Data were collected for 2017–2025 to align with available mosquito surveillance data from Richardson, Lancaster, and Douglas Counties. Data retrieval and processing were conducted in R using the httr2 (Wickham 2025a), lubridate (Grolemund and Wickham 2011), and jsonlite (Ooms 2014) packages to access and parse API responses (Supp. Fig. S1).

### Degree-Day Accumulation and Emergence Timing Assessment

Cumulative degree-days were calculated daily beginning January 1 of each year. Adult emergence was predicted to occur when cumulative thermal accumulation reached or exceeded the developmental threshold of 172.4 CDD (Healy et al. 2019). The earliest date within each year on which this threshold was met or exceeded was designated as the predicted emergence date. Observed emergence was approximated using surveillance data. For each year, the first detection was defined as the earliest collection date on which at least one *Ae. albopictus* individual was recorded following the initiation of trapping.

To visualize emergence phenology, cumulative degree-day trajectories were plotted across the sampling season for each year. Vertical reference lines were used to indicate both predicted emergence dates based on CDD thresholds and first detection dates from surveillance data. Differences between predicted and observed timing were interpreted as indicators of model performance, as well as potential environmental or sampling-related influences on mosquito phenology. Graphs were produced using the ggplot2 and ggbreak packages (v4.0.3; Wickham 2016, Xu et al. 2021).

First detection in surveillance collections was used as a proxy for adult emergence because direct measurement of emergence from natural larval habitats is not feasible at landscape scales. Standardized trapping methods instead capture host-seeking or dispersing adults, providing the most practical field-based estimate of emergence timing.

### Statistical analysis

All analyses were conducted in RStudio (version 4.3.1) using reproducible, script-based workflows (R Core Team 2023). Temperature data were accessed via the NASA POWER API, and degree-day calculations were implemented using custom R functions (NASA LaRC 2026). Data management was completed in RStudio using dplyr (Wickham et al. 2026a), tibble (Müller and Wickham 2026), stringr (Wickham 2025b), and readr (Wickham et al. 2026b) (Supp. Fig. S1).

To evaluate whether urban environments influenced the probability of *Ae. albopictus* detection, we modeled annual trap positivity as a function of trap location relative to municipal boundaries. Trap locations were classified as either inside or outside city limits using spatial intersections with U.S. Census place boundaries for Lincoln (Lancaster County), Omaha (Douglas County), and Falls City (Richardson County).

For each trap location and year (2017–2025), detection was coded as a binary response (positive = ≥1 *Ae. albopictus* detected; negative = none detected). Because traps were sampled repeatedly across years, generalized linear mixed-effects models (GLMMs) with a binomial error distribution and logit link were used, with trap identity included as a random intercept to account for repeated measures.

Due to differences in invasion stage among counties and evidence of quasi-complete separation in pooled analyses, models were fit separately by county. For Douglas and Lancaster Counties, GLMMs were implemented using the lme4 (Bates et al. 2015) and pracma (Borchers 2025) packages. For Richardson County, where separation was observed, penalized logistic regression with Firth correction was used via the brglm2 package to obtain bias-reduced parameter estimates (Kosmidis 2026).

## Results

### Surveillance and detection of Ae. albopictus in Lincoln, NE

Surveillance was conducted in 44 counties across Nebraska during 2017–2025 (Fig. 1A), with 14 counties sampled consistently throughout all nine years (Fig. 1A). Over this period, a total of 1,145,412 mosquitoes were collected, of which 6,093 (0.53%) were identified as *Ae. albopictus* (Fig. 1C). These data indicate an emerging but spatially heterogeneous distribution across counties.

**Figure 1.**
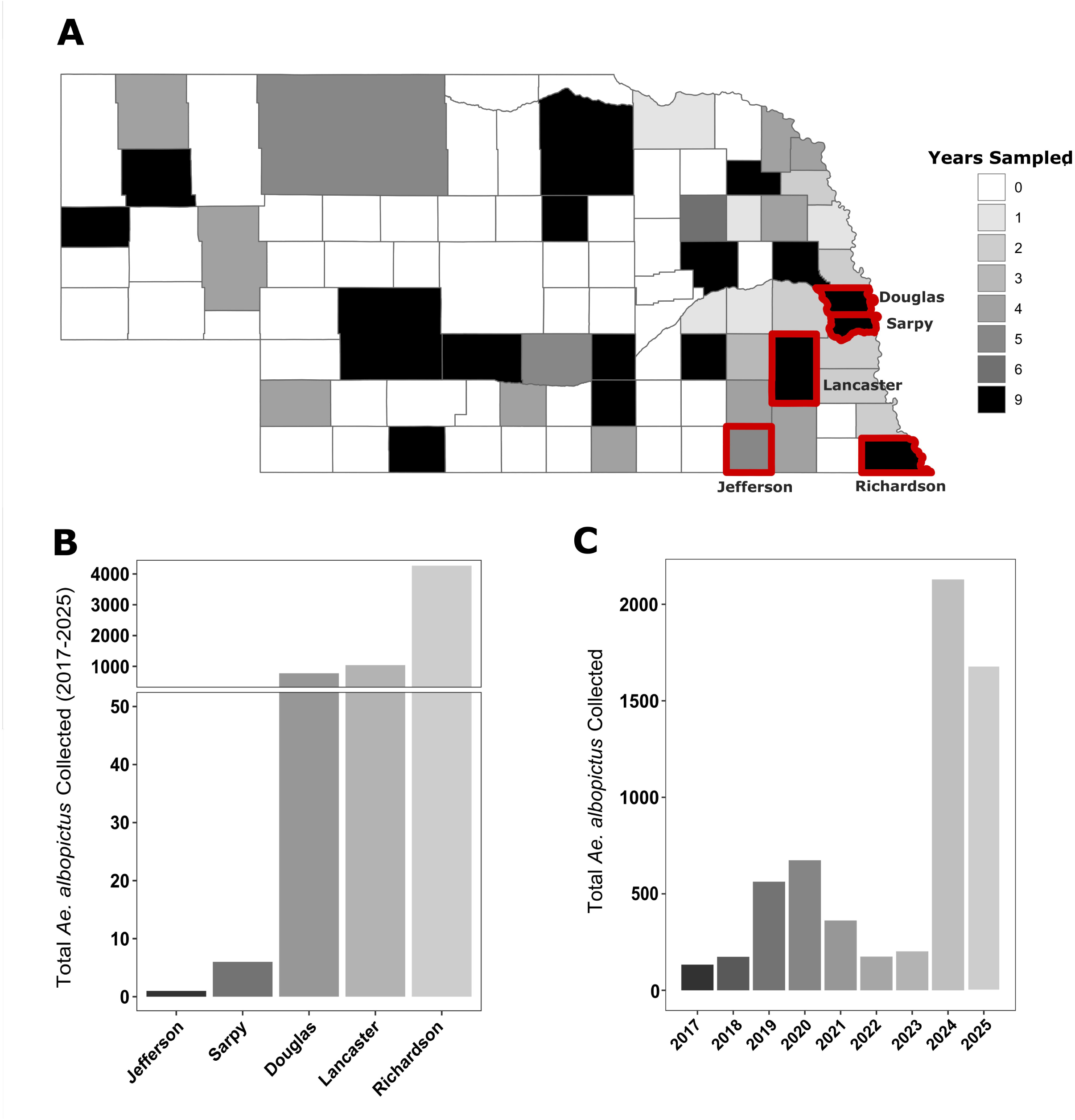
Overview of mosquito trapping locations and the number of *Ae. albopictus* collected in Nebraska during 2017–2025. Participating counties are shaded based on the number of years trapping was conducted (**A**). Counties in which *Ae. albopictus* was detected are outlined in red. The number of *Ae. albopictus* collected is displayed for each positive county (**B**) and for each year (**C).**

Richardson County exhibited the earliest and most consistent detection of *Ae. albopictus*. The species was first detected in 2017, comprising 4.56% of the total mosquitoes collected, and was recorded annually thereafter. Relative abundance increased from 5.88% in 2018 to 10.39% in 2019 and 40.28% in 2020, followed by a peak of 56.41% in 2021. Moderate declines were observed in 2022 (27.53%) and 2023 (24.35%), with proportions stabilizing in 2024 (27.27%) before reaching the highest recorded value of 60.50% in 2025. Across the study period, Richardson County accounted for 4,267 total (70.0%) *Ae. albopictus* specimens (Fig. 1B; Table 1).

**Table 1.** Percentage of *Ae. albopictus* identified out of the total mosquitoes collected in Douglas, Jefferson, Lancaster, Richardson, and Sarpy counties (2017–2025).

| County | 2017 | 2018 | 2019 | 2020 | 2021 | 2022 | 2023 | 2024 | 2025 |
| --- | --- | --- | --- | --- | --- | --- | --- | --- | --- |
| Douglas | 0 | 0.005 | 0.007 | 0.023 | 0.016 | 0.69 | 2.33 | 5.22 | 0.78 |
| Jefferson | 0 | 0 | 0 | 0 | 0 | 0.11 | NA | NA | NA |
| Lancaster | 0 | 0 | 0 | 0 | 0 | 0.27 | 0.52 | 6.06 | 6.87 |
| Richardson | 4.56 | 5.88 | 10.39 | 40.28 | 56.41 | 27.53 | 24.35 | 27.27 | 60.5 |
| Sarpy | 0 | 0 | 0 | 0 | 0 | 0 | 0 | 0.061 | 0.062 |

Douglas County represented a later phase of detection. *Ae. albopictus* was first recorded in 2018, comprising only 0.005% of total mosquito collections. Relative abundance remained very low through 2019 (0.007%), 2020 (0.023%), and 2021 (0.016%). A notable increase was observed in 2022 (0.69%), followed by continued growth in 2023 (2.33%), reaching a peak of 5.22% in 2024 before declining to 0.78% in 2025. Across all sampling years, a total of 778 total (12.9%) *Ae. albopictus* were collected in Douglas County (Fig. 1B; Table 1).

Lancaster County exhibited the latest detection timeline among the primary counties. The first confirmed presence occurred in 2022, when *Ae. albopictus* comprised 0.27% of total mosquito collections. The proportion increased in 2023 (0.52%), followed by substantial growth in 2024 (6.06%) and 2025 (6.87%). Over the study period, Lancaster County accounted for 1,041 total (17.1%) *Ae. albopictus* (Fig. 1B; Table 1).

Detections outside of these three counties were limited. In Sarpy County, *Ae. albopictus* was recorded in 2024 (0.061%) and 2025 (0.062%), totaling six individuals (Fig. 1B; Table 1). In Jefferson County, a single *Ae. albopictus* was recorded in 2022, representing 0.11% of that year’s collections, with no subsequent detections (Fig. 1B; Table 1).

### Cumulative degree-day modeling

Degree-day modeling and trap persistence analyses were restricted to Richardson, Douglas, and Lancaster counties due to sufficient longitudinal coverage and repeated detection of *Ae. albopictus*. Sarpy and Jefferson counties were excluded because detections were sporadic and sample sizes were insufficient to support robust temporal or spatial analyses.

Across all three counties and all sampled years, the CDD threshold of 172.4 was consistently reached between May 8 and May 22 (Table 2; Figures 2–4). In every county and year, initial trap detection occurred after the predicted emergence date. This lag ranged from approximately two to four weeks in Lancaster and Douglas counties and was more variable in Richardson County, spanning from nine days to more than seven weeks. The most extreme delay occurred in Richardson County in 2021, when traps were not deployed until July 13, corresponding to a CDD of 884.01, which far exceeds the 172.4 threshold and is likely due to COVID-19-related disruptions and funding constraints.

**Table 2.** Dates and cumulative degree days (CDD) for predicted collection, first trapping, and first *Ae. Albopictus* collection.

| County | Predicted Collection Date | Predicted Collection CDD | First Trapping Date | First Trapping CDD | *Ae. albopictus* First Collection Date | *Ae. albopictus* First Collection CDD |
| --- | --- | --- | --- | --- | --- | --- |
| Lancaster | 16-5-2022 | 172.4 | 9-6-2022 | 369.27 | 19-7-2022 | 998.67 |
| Lancaster | 13-5-2023 | 172.4 | 9-6-2023 | 476.57 | 15-8-2023 | 1431.23 |
| Lancaster | 16-5-2024 | 172.4 | 4-6-2024 | 353.74 | 4-6-2024 | 353.74 |
| Lancaster | 10-5-2025 | 172.4 | 5-6-2025 | 372.29 | 10-6-2025 | 417.8 |
| Douglas | 15-5-2022 | 172.4 | 1-6-2022 | 279.31 | 23-8-2022 | 1557.96 |
| Douglas | 13-5-2023 | 172.4 | 31-5-2023 | 328.78 | 27-7-2023 | 1118.05 |
| Douglas | 15-5-2024 | 172.4 | 29-5-2024 | 277.7 | 29-5-2024 | 277.7 |
| Douglas | 11-5-2025 | 172.4 | 28-5-2025 | 266.9 | 8-7-2025 | 811.96 |
| Richardson | 17-5-2017 | 172.4 | 31-5-2017 | 329.15 | 31-5-2017 | 329.15 |
| Richardson | 12-5-2018 | 172.4 | 29-5-2018 | 404.87 | 30-6-2018 | 628.18 |
| Richardson | 22-5-2019 | 172.4 | 6-6-2019 | 327.23 | 6-6-2019 | 327.23 |
| Richardson | 22-5-2020 | 172.4 | 27-5-2020 | 220.18 | 8-6-2020 | 371.24 |
| Richardson | 19-5-2021 | 172.4 | 13-7-2021 | 884.01 | 10-8-2021 | 1329.38 |
| Richardson | 13-5-2022 | 172.4 | 3-6-2022 | 346.12 | 3-6-2022 | 346.12 |
| Richardson | 13-5-2023 | 172.4 | 1-6-2023 | 349.48 | 1-6-2023 | 349.48 |
| Richardson | 8-5-2024 | 172.4 | 30-5-2024 | 372.7 | 30-5-2024 | 372.7 |
| Richardson | 8-5-2025 | 172.4 | 29-5-2025 | 332.46 | 29-5-2025 | 332.46 |

**Figure 2.**
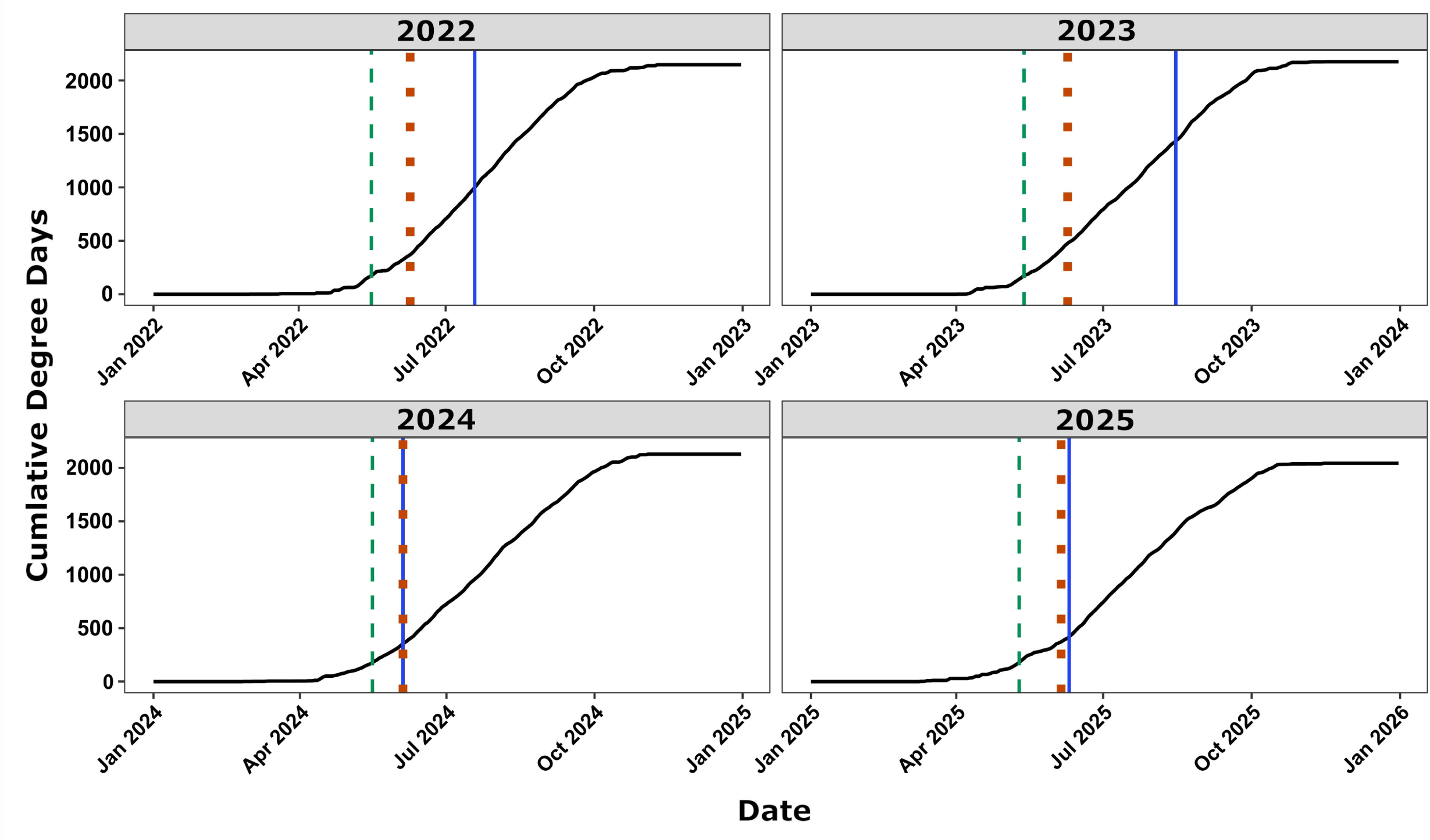
Lancaster County cumulative degree days during 2022–2025. The Green dashed line denotes the predicted emergence, the orange dotted line indicates the first trap collection, and the blue line represents the first observed *Ae. albopictus* collected in the traps.

**Figure 3.**
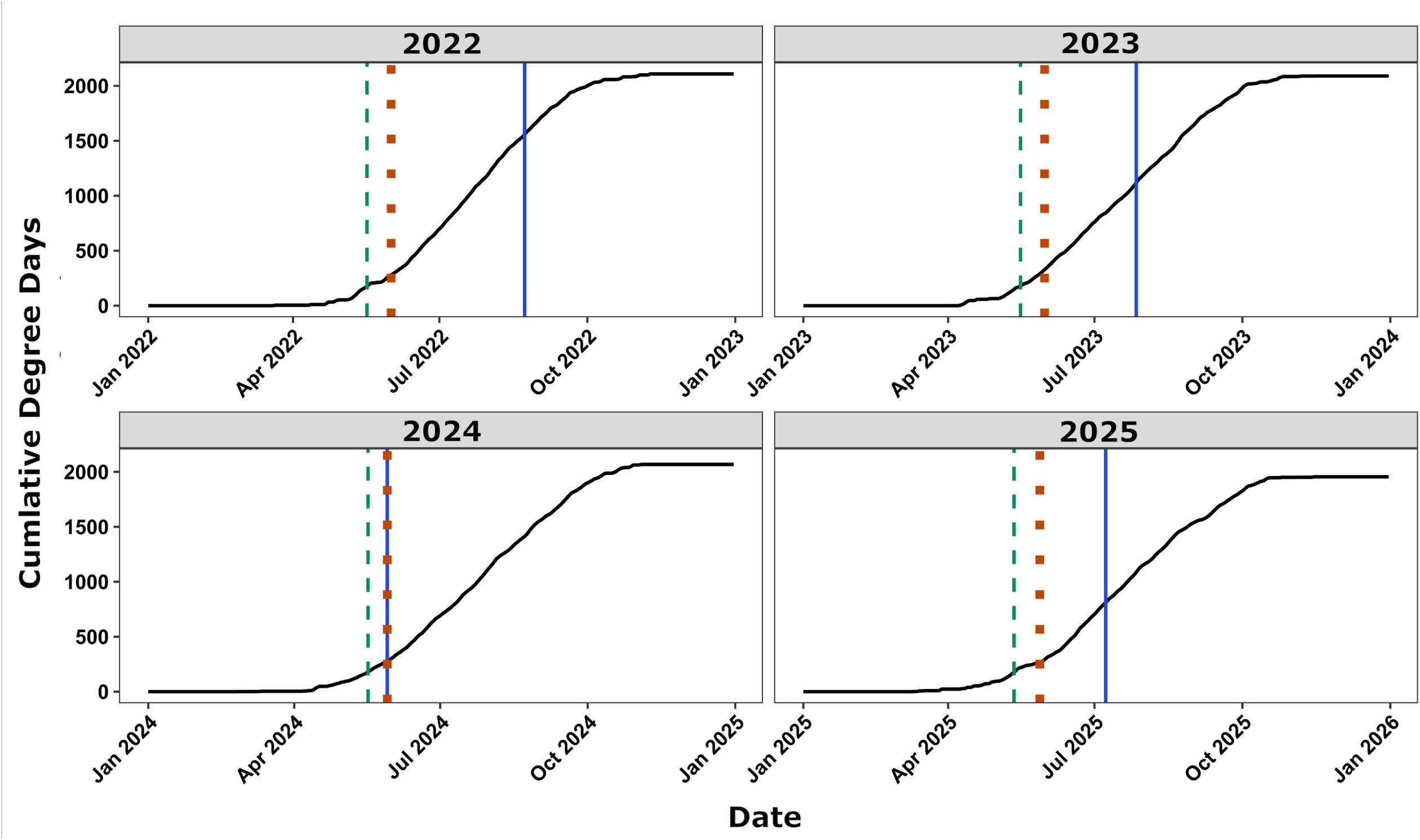
Douglas County cumulative degree days during 2022–2025. The green dashed line denotes the predicted emergence, the orange dotted line indicates the first trap collection, and the blue line represents the first observed *Ae. albopictus* collected in the traps.

**Figure 4.**
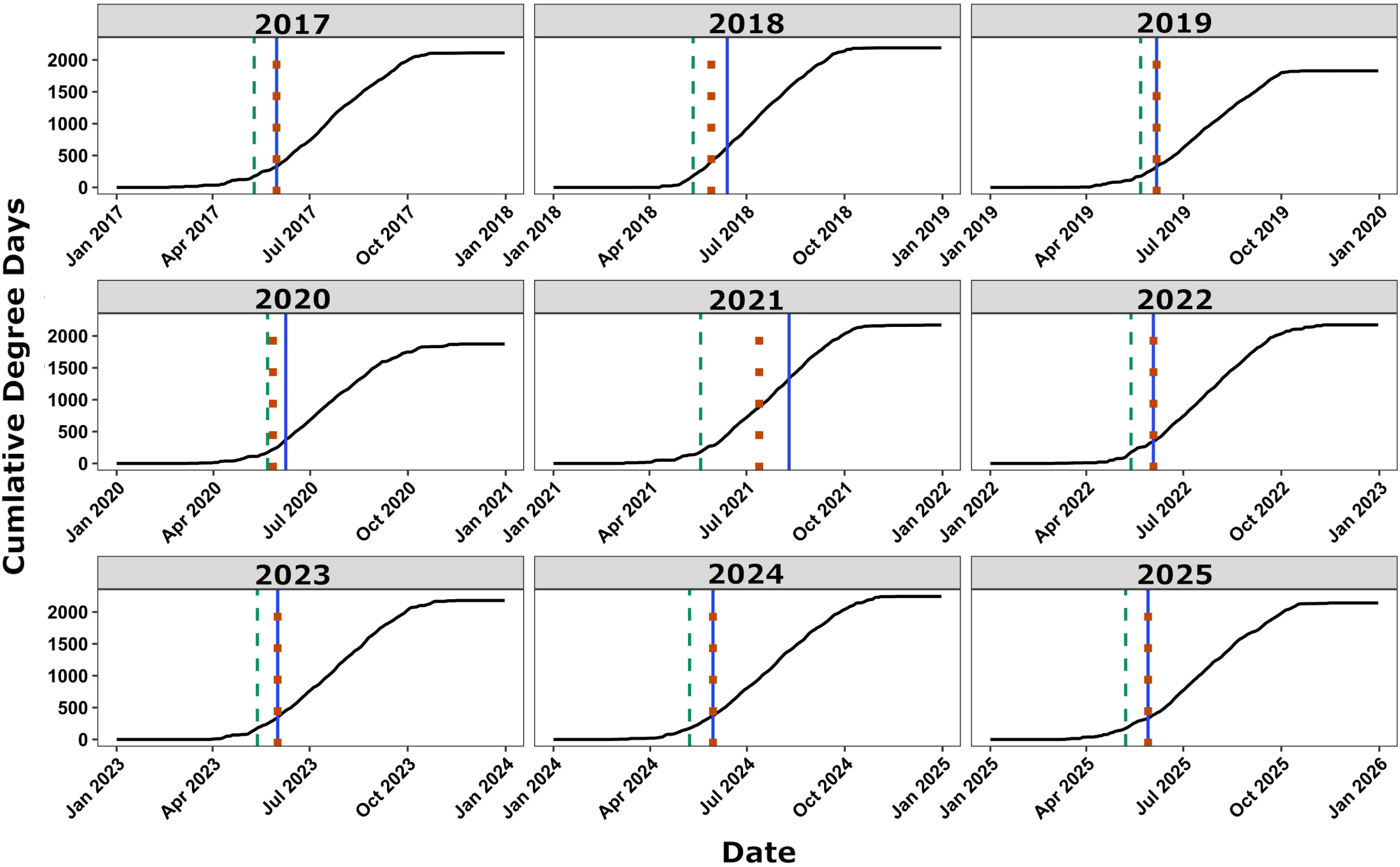
Richardson County cumulative degree days during 2017–2025. The green dashed line denotes the predicted emergence, the orange dotted line indicates the first trap collection, and the blue line represents the first observed *Ae. albopictus* collected in the traps.

A consistent pattern was observed across all three counties: in multiple years, *Ae. albopictus* was detected on the first day of trap deployment. This occurred in Richardson County in 2017, 2019, 2023, 2024, and 2025, and in both Lancaster and Douglas counties in 2024 (Table 2; Figures 2–4). In these cases, the CDD at first detection was identical to that at trap deployment.

In Lancaster and Douglas counties, earlier trap deployment relative to the predicted emergence date was associated with earlier detection of *Ae. albopictus*, whereas delayed trap deployment corresponded to higher CDD values at first detection. For example, in Douglas County, the CDD at first detection ranged from 277.7 in 2024 to 1,557.96 in 2022, reflecting a longer interval between trap deployment and detection in the latter year. A similar pattern was observed in Lancaster County, where first detection occurred at 998.67 CDD in 2022 and 1,431.23 CDD in 2023, compared to 353.74 CDD in 2024 when detection coincided with initial trap deployment.

Richardson County, which has the longest surveillance record (2017–2025), showed the most consistent pattern of early detection relative to trap deployment, with same-day detection occurring in five of nine years.

### Urban effect on the establishment of Ae. albopictus

Across counties, *Ae. albopictus* detections were highly spatially clustered, with repeated positives concentrated at a limited subset of trap locations (Fig. 5). Although raw detection frequencies were higher within city limits than outside, generalized linear mixed-effects models indicated that urban location was not a significant predictor of yearly trap positivity after accounting for repeated sampling at individual trap locations. In Douglas County, the effect of urban location was positive but not significant (*β = 1.53, SE = 3.57, z = 0.43, P = 0.67; OR = 4.63, 95% CI: 0.004–5046*) (Fig. 5B). Similarly, in Lancaster County, urban location showed a positive but non-significant association with detection (*β = 3.28, SE = 7.02, z = 0.47, P = 0.64; OR = 26.7, 95% CI: 2.8 × 10⁻⁵–2.5 × 10⁷*) (Fig. 5A).

**Figure 5.**
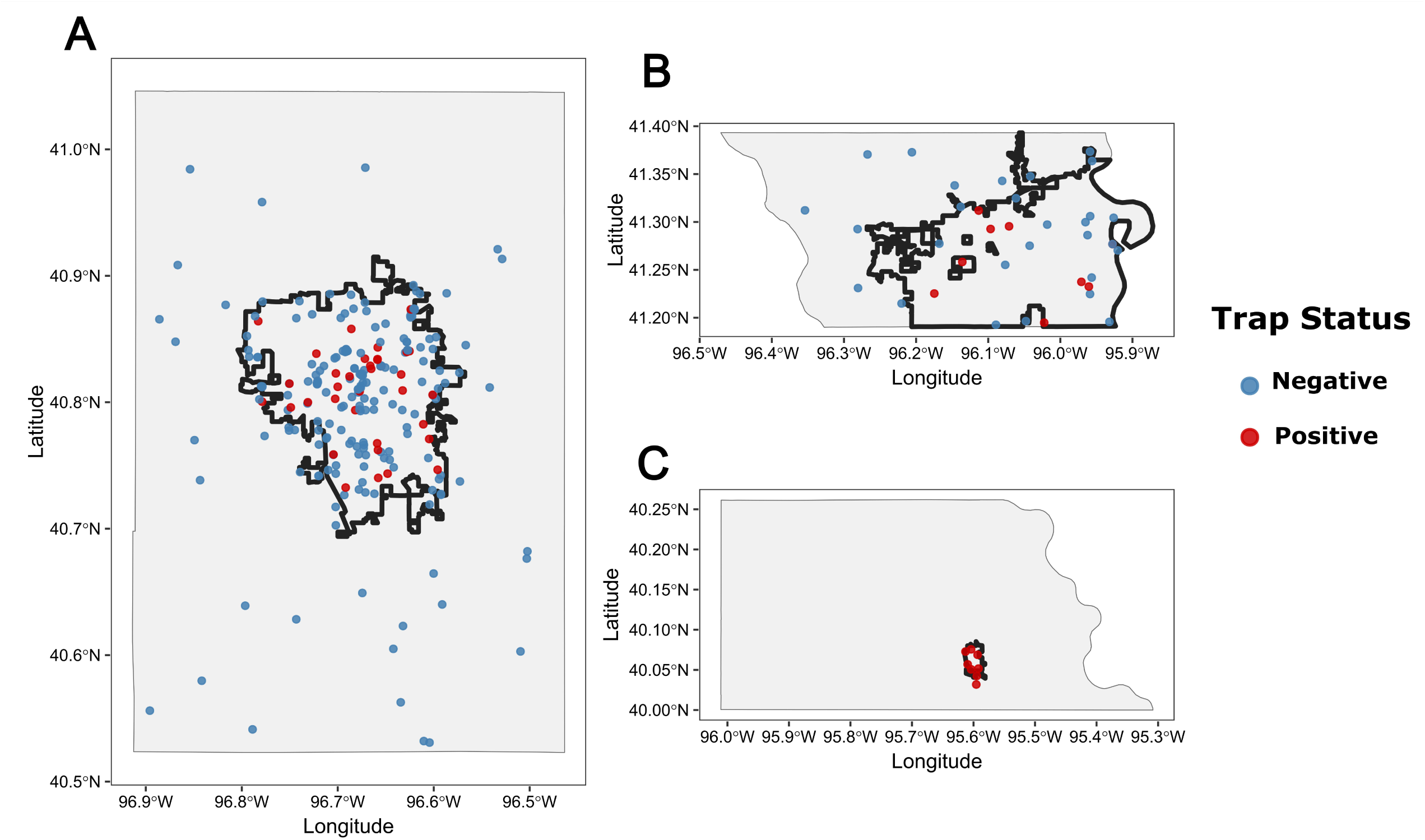
Trap locations in Lancaster (A), Douglas (B), and Richardson (C) during 2017–2025. City limits are outlined with the solid black line for Lincoln (A), Omaha (B), and Falls City (C). Traps that were positive for *Ae. albopictus* are highlighted in red, and traps that never detected *Ae. albopictus* are highlighted in blue.

In contrast, trap-level persistence was a strong predictor of detection probability across the full dataset. Traps with higher persistence, defined as the proportion of sampled years in which *Ae. albopictus* was detected, had substantially higher odds of yearly positivity (*β = 8.98, SE = 0.93, z = 9.69, P < 0.001*). Inclusion of persistence reduced the random intercept variance for trap identity to approximately 0, indicating that persistent, site-specific differences accounted for nearly all observed heterogeneity in detection probability.

## Discussion

Surveillance conducted across 44 Nebraska counties during 2017–2025 documents the progressive establishment of *Ae. albopictus* in eastern Nebraska. Sustained detections in Richardson County, along with emerging establishments in Lancaster and Douglas counties beginning in 2022, indicate continued expansion within the region. These findings are consistent with the broader pattern of northward and westward range expansion reported across the central and eastern U.S. (Hahn et al. 2017) and extend the northern limit of confirmed *Ae. albopictus* populations in the Great Plains. The detection patterns observed across counties follows an invasion trajectory characterized by initial colonization followed by progressive local population growth, consistent with patterns documented in neighboring states such as Iowa and Missouri (Claborn et al. 2018, Hall et al. 2022).

Richardson County provides the strongest evidence of establishment, with continuous annual detections during 2017–2025 and a sustained increase in proportional representation, reaching 60.50% of collected mosquitoes in 2025. This trajectory is unlikely to reflect repeated independent introductions and instead supports the presence of a self-sustaining population undergoing local growth. The pattern closely mirrors establishment dynamics reported in Iowa, where progressive increases in detection frequency, coupled with genetic evidence, indicated the development of a stable breeding population following initial colonization (Hall et al. 2022). In Douglas and Lancaster counties, the most compelling evidence for establishment arises not from initial detection but from the temporal advancement of seasonal activity in 2024 and 2025. In these years, *Ae. albopictus* was detected earlier in the trapping season and at substantially lower CDD values compared to 2022 and 2023. This pattern is inconsistent with an annual reintroduction hypothesis, as passive dispersal events would not be expected to produce a consistent, multi-year shift toward earlier seasonal detection (Medlock et al. 2012, Lyberger et al. 2025). Instead, earlier seasonal activity is consistent with successful overwintering of diapausing eggs, enabling local populations to resume development from an established egg bank in spring rather than requiring *de novo* introduction each year (Nawrocki and Hawley 1987, Lee et al. 2024).

A key challenge in interpreting these findings is the absence of a universally accepted operational definition of *Ae. albopictus* establishment, particularly in distinguishing self-sustaining local populations from recurrent seasonal introductions (Hall et al. 2022, Lee et al. 2026). In this study, establishment is defined as the repeated detection of *Ae. albopictus* at the same trap locations across multiple consecutive years, accompanied by evidence of increasing population abundance and earlier seasonal activity. Although this definition is practical and consistent with prior surveillance studies (Hall et al. 2022, Lee et al. 2026), it does not provide direct biological confirmation of successful overwintering in the absence of egg or larval data. Future efforts incorporating oviposition traps deployed during late fall through early spring, or larval sampling from known container habitats, would offer stronger evidence for population persistence. Establishing a standardized and broadly accepted framework for defining *Ae. albopictus* establishment would strengthen vector surveillance efforts, particularly as the species continues expanding into previously unoccupied regions of the northern U.S.

The absence of a statistically significant urban effect on detection probability, after accounting for trap-level persistence, has important implications for surveillance design and invasion risk assessment. GLMM results indicate that the concentration of detections within city limits is driven primarily by consistent positivity at a limited number of trap locations, rather than by a broad urban effect. This pattern is consistent with invasion theory, which predicts that early-stage establishment is governed by localized propagule pressure and microhabitat suitability at specific introduction points, rather than by landscape-scale factors such as urbanization (Lockwood et al. 2005, Blackburn et al. 2014). Although urban environments may facilitate initial colonization through increased human connectivity, greater availability of artificial container habitats, and elevated temperatures associated with the urban heat island effect (Medlock et al. 2012, Wiese et al. 2019), these influences appear secondary to site-specific ecological conditions during early invasion. As populations expand and the number of source locations increase, urbanization may emerge as a stronger predictor of detection probability, as observed in more advanced invasion fronts in the eastern U.S. (Rochlin et al. 2013).

The cumulative degree-day model of Healy et al. (2019) consistently predicted adult *Ae. albopictus* emergence in mid-May across all three counties and all years examined, yet trap deployment by the health departments did not begin until after this predicted emergence window in every county-year combination, creating a systematic surveillance gap of approximately two to seven weeks. This lag is largely attributable to the broader focus of the DHHS trapping program on overall mosquito abundance, species richness, and virus surveillance for *Culex,* rather than targeted *Ae. albopictus* phenology monitoring. As a result, surveillance-derived estimates of first detection date, seasonal abundance, and population phenology are biased toward later dates and likely underestimate true abundance. Aligning trap deployment with model-predicted emergence could substantially improve surveillance fidelity and enable more proactive vector control, targeting early-season adults before populations increase (Fonseca et al. 2013, Armstrong et al. 2017, Healy et al. 2019).

Several limitations should be considered when interpreting these results. Sampling effort varied across counties and years, aligned with personnel availability, logistical constraints, and during 2020 and 2021, resource reallocation associated with the COVID-19 pandemic and the loss of prior Zika-related funding. Additionally, CO₂-baited CDC light traps, the primary trapping tool used throughout the program, are known to be less effective at capturing container-breeding *Aedes* species than host-mimicking traps such as the BG-Sentinel (Maciel-de-Freitas et al. 2006, Farajollahi et al. 2009), meaning *Ae. albopictus* abundance and distribution were likely underestimated in years and locations where only CDC light traps were deployed. Despite these constraints, the progressive establishment of *Ae. albopictus* in eastern Nebraska has meaningful public health implications. As a competent vector of West Nile virus with the capacity for transovarial transmission, *Ae. albopictus* could contribute to pathogen persistence through winter diapause and introduce a secondary transmission pathway supplementing the existing *Culex*-dominated enzootic cycle (Zhang et al. 2022). The preliminary detections in Sarpy County in 2024 and 2025 suggest the invasion front continues to expand. Integrating degree-day model predictions into existing health department surveillance schedules represents a practical, evidence-based approach to improving the timeliness and sensitivity of *Ae. albopictus* monitoring in Nebraska.

This study documents the detection and likely establishment of *Ae. albopictus* in three Nebraska counties over a nine-year surveillance period, representing the northernmost confirmed presence of the species in the central Great Plains. Richardson County showed continuous annual detections beginning in 2017, with proportional abundance increasing steadily to more than 60% of collected mosquitoes by 2025. In Douglas and Lancaster counties, the earlier timing of first seasonal detection in 2024 and 2025, relative to previous years, is consistent with successful overwintering rather than repeated reintroduction. Across all three counties, the cumulative degree-day model of Healy et al. (2019) consistently predicted adult emergence in mid-May, preceding trap deployment by two to seven weeks and highlighting a persistent early-season surveillance gap. Trap-level persistence, rather than urban proximity, was the strongest predictor of yearly *Ae. albopictus* detection, suggesting that current invasion dynamics in Nebraska are driven by localized source populations rather than broad landscape-scale urbanization. As *Ae. albopictus* continues to expand northward and westward, earlier trap deployment aligned with degree-day model predictions, together with oviposition or larval monitoring to confirm overwintering, will be critical for accurately characterizing population phenology and supporting timely vector control in Nebraska and similar continental climate regions.

## Acknowledgments

The temperature data used in this study was obtained from the National Aeronautics and Space Administration (NASA) Langley Research Center (LaRC) Prediction of Worldwide Energy Resource (POWER) Project, which is funded through the NASA Applied Sciences Program within the Earth Science Division of the Science Mission Directorate.

