## Supplemental Figure 1 for "Long-Term Surveillance Reveals Establishment of *Aedes albopictus* in Eastern Nebraska, USA": Supplemental_Figure_1.html

Supplemental Figure S1


### Supplemental Figure S1

###### Gwenyth Coleman

#### 2026-09-16

#### Code for Cumulative Degree-Day (CDD) Modeling of Lancaster, Douglas and Richardson counties.

Loading Libraries

```
library(httr) # (Wickham 2025a)
library(jsonlite) # (Ooms 2014)
library(dplyr) # (Wickham et al. 2026a)
library(lubridate) # (Grolemund and Wickham 2011)
library(tibble) # (Müller and Wickham 2026)
library(stringr) #(Wickham 2025b)
library(pracma) # (Borchers 2025)
library(ggplot2) # (v4.0.3; Wickham 2016)
library(readr) # (Wickham et al. 2026b)
```

Loading and filtering the raw data for *Ae. albopictus*

```
#Lancaster County
mosq_lan <- read_csv("./Supplemental_Table_1.csv") %>%
  mutate(
    date = mdy(collection_date),
    year = year(date),
    county = str_to_title(calculated_county),
    alb = coalesce(aedes_albopictus_males, 0) +
      coalesce(`aedes_albopictus_females-mixed`, 0)
  ) %>%
  filter(county == "Lancaster")

# Douglas County
mosq_dug <- read_csv("./Supplemental_Table_1.csv") %>%
  mutate(
    date = mdy(collection_date),
    year = year(date),
    county = str_to_title(calculated_county),
    alb = coalesce(aedes_albopictus_males, 0) +
      coalesce(`aedes_albopictus_females-mixed`, 0)
  ) %>%
  filter(county == "Douglas")

#Richardson County
mosq_ric <- read_csv("./Supplemental_Table_1.csv") %>%
  mutate(
    date = mdy(collection_date),
    year = year(date),
    county = str_to_title(calculated_county),
    alb = coalesce(aedes_albopictus_males, 0) +
      coalesce(`aedes_albopictus_females-mixed`, 0)
  ) %>%
  filter(county == "Richardson")
```

Development temperatures and cumulative degree days taken from Healy
et al. (2019) using the lab population

```
Tmin  <- 10.5
Tpeak <- 32.93
Tmax  <- 36.2
CDD_final <- 172.4
```

Function to collect weather data for the counties from National
Aeronautics and Space Administration (NASA) Langley Research Center
(LaRC) Prediction of Worldide Energy (POWER) (NASA LaRC 2026)

```
get_nasa_power = function(lat, lon, start_date, end_date) {
  
  base_url = "https://power.larc.nasa.gov/api/temporal/daily/point"
  
  response <- GET(
    url = base_url,
    query = list(
      start = gsub("-", "", start_date),
      end   = gsub("-", "", end_date),
      latitude = lat,
      longitude = lon,
      community = "AG",
      parameters = "T2M,T2M_MIN,T2M_MAX",
      format = "JSON"
    )
  )
  
  data_json = content(response, "text")
  data = fromJSON(data_json)
  
  df = data$properties$parameter
  
  tibble(
    date = seq(as.Date(start_date), as.Date(end_date), by = "day"),
    Tmean = as.numeric(df$T2M),
    Tmin  = as.numeric(df$T2M_MIN),
    Tmax  = as.numeric(df$T2M_MAX)
  ) %>%
    filter(!is.na(Tmean))  # Remove days without data
}
```

Temperature data acquisition from Lancaster, Douglas, and Richardson
county from 2017-2025 (NASA LaRC 2026)

```
#Lancaster county
temp_lancaster <- get_nasa_power(
  lat = 40.8471,
  lon = -96.6638,
  start_date = "2022-01-01",
  end_date   = "2025-12-31"
)

#Douglas county
temp_douglas <- get_nasa_power(
  lat = 41.31,
  lon = -96.2,
  start_date = "2022-01-01",
  end_date   = "2025-12-31"
)

#Richardson county
temp_richardson <- get_nasa_power(
  lat = 40.0833,
  lon = -95.7000,
  start_date = "2017-01-01", 
  end_date   = "2025-12-31"
)
```

Creation of a function that calculates CDD using a nonlinear
temperature-response function based on mean daily temperature (Healy et
al. 2019)

```
calc_dd_nonlinear <- function(Tmean, Tmin = 10.7, Tpeak = 32.35, Tmax = 36.8) {
  if (Tmean <= Tmin) return(0)
  if (Tmean >= Tpeak & Tmean <= Tmax) return(Tpeak - Tmin)
  if (Tmean > Tmax) return(0)
  return(Tmean - Tmin)
}
```

Using the function calculating CDD for each county

```
#Lancaster County
dd_lancaster <- temp_lancaster %>%
  mutate(
    dd = sapply(Tmean, calc_dd_nonlinear),
    year = year(date),
    doy = yday(date),
    CDD = ave(dd, year, FUN = cumsum)
  )

# Douglas County
dd_douglas <- temp_douglas %>%
  mutate(
    dd = sapply(Tmean, calc_dd_nonlinear),
    year = year(date),
    doy = yday(date),
    CDD = ave(dd, year, FUN = cumsum)
  )

# Richarson County
dd_richardson <- temp_richardson %>%
  mutate(
    dd = sapply(Tmean, calc_dd_nonlinear), #degree days
    year = year(date),
    doy = yday(date),
    CDD = ave(dd, year, FUN = cumsum) #cumulative degree days for the years
  )
```

Calculating mosquito counts per county

```
#Lancaster County
mosq_rich_lan <- mosq_lan %>%
  filter(county == "Lancaster") %>%
  arrange(date) %>%
  mutate(year = year(date)) %>%
  group_by(year) %>%
  mutate(
    total = sum(alb, na.rm = TRUE),
    cum = cumsum(alb),
    prop = ifelse(total > 0, cum / total, NA_real_)
  ) %>%
  ungroup()

#Douglas County
mosq_rich_dug <- mosq_dug %>%
  filter(county == "Lancaster") %>%
  arrange(date) %>%
  mutate(year = year(date)) %>%
  group_by(year) %>%
  mutate(
    total = sum(alb, na.rm = TRUE),
    cum = cumsum(alb),
    prop = ifelse(total > 0, cum / total, NA_real_)
  ) %>%
  ungroup()

#Richardson County
mosq_rich_ric <- mosq_ric %>%
  filter(county == "Lancaster") %>%
  arrange(date) %>%
  mutate(year = year(date)) %>%
  group_by(year) %>%
  mutate(
    total = sum(alb, na.rm = TRUE),
    cum = cumsum(alb),
    prop = ifelse(total > 0, cum / total, NA_real_)
  ) %>%
  ungroup()
```

Identify emergence year

```
#Lancaster County
first_emergence_lan <- mosq_rich_lan %>%
  filter(alb > 0) %>%
  group_by(year) %>%
  summarise(
    first_emergence_date_lan = min(date),
    .groups = "drop"
  )

#Douglas County
first_emergence_dug <- mosq_rich_dug %>%
  filter(alb > 0) %>%
  group_by(year) %>%
  summarise(
    first_emergence_date_dug = min(date),
    .groups = "drop"
  )

#Richardson County
first_emergence_ric <- mosq_rich_ric %>%
  filter(alb > 0) %>%
  group_by(year) %>%
  summarise(
    first_emergence_date_ric = min(date),
    .groups = "drop"
  )
```

Identify date when CDD reaches the threshold

```
#Lancaster County
dd_threshold_lan <- dd_lancaster %>%
  group_by(year) %>%
  filter(CDD >= CDD_final) %>%
  summarise(
    dd_threshold_date_lan = min(date),
    .groups = "drop"
  )

#Douglas County
dd_threshold_dug <- dd_douglas %>%
  group_by(year) %>%
  filter(CDD >= CDD_final) %>%
  summarise(
    dd_threshold_date_dug = min(date),
    .groups = "drop"
  )

#Richardson County
dd_threshold_ric <- dd_richardson %>%
  group_by(year) %>%
  filter(CDD >= CDD_final) %>%
  summarise(
    dd_threshold_date_ric = min(date),
    .groups = "drop"
  )
```

Identify first trap collection of any species per year and joining
data for plotting

```
#Lancaster
first_collection_overall_lan <- mosq_lan %>%
  group_by(year) %>%
  summarise(
    first_collection_date_lan = min(date),
    .groups = "drop"
  )

dd_plot_data_lan <- dd_lancaster %>%
  left_join(first_emergence_lan, by = "year") %>%
  left_join(first_collection_overall_lan, by = "year") %>%
  left_join(dd_threshold_lan, by = "year")

#Douglas County
first_collection_overall_dug <- mosq_dug %>%
  group_by(year) %>%
  summarise(
    first_collection_date_dug = min(date),
    .groups = "drop"
  )

dd_plot_data_dug <- dd_douglas %>%
  left_join(first_emergence_dug, by = "year") %>%
  left_join(first_collection_overall_dug, by = "year") %>%
  left_join(dd_threshold_dug, by = "year")

#Richardson County
first_collection_overall_ric <- mosq_ric %>%
  group_by(year) %>%
  summarise(
    first_collection_date_ric = min(date),
    .groups = "drop"
  )

dd_plot_data_ric <- dd_richardson %>%
  left_join(first_emergence_ric, by = "year") %>%
  left_join(first_collection_overall_ric, by = "year") %>%
  left_join(dd_threshold_ric, by = "year")
```

#### Plotting CDD

Lancaster County

```
ggplot(dd_plot_data_lan, aes(x = date, y = CDD)) +
  geom_line(color = "black", linewidth = 0.8) +
  
  # DD threshold line
  geom_vline(
    aes(xintercept = dd_threshold_date_lan),
    linetype = "dashed",
    color = "#029356",
    linewidth = 0.8
  ) +
  
  # First emergence line
  geom_vline(
    aes(xintercept = first_emergence_date_lan),
    linetype = "solid",
    color = "#2546f0",
    linewidth = 0.8
  ) + 
  # First sampling
  geom_vline(
    aes(xintercept = first_collection_date_lan),
    linetype = "dotted",
    color = "#c44601",
    linewidth = 2
  ) +
  
  facet_wrap(~year, scales = "free_x") +
  
  theme_bw() +
  theme(
    strip.text = element_text(face = "bold"),
    axis.text.x = element_text(face = "bold", angle = 45, hjust = 1, , size = 10),
    axis.text.y = element_text(face = "bold", size = 10),
    panel.grid.major = element_blank(),
    panel.grid.minor = element_blank()
  ) +
  
  labs(
    x = "Date",
    y = "Cumulative Degree Days"
  )
```

Douglas County

```
ggplot(dd_plot_data_dug, aes(x = date, y = CDD)) +
  geom_line(color = "black", linewidth = 0.8) +
  
  # DD threshold line
  geom_vline(
    aes(xintercept = dd_threshold_date_dug),
    linetype = "dashed",
    color = "#029356",
    linewidth = 0.8
  ) +
  
  # First emergence line
  geom_vline(
    aes(xintercept = first_emergence_date_dug),
    linetype = "solid",
    color = "#2546f0",
    linewidth = 0.8
  ) + 
  # First sampling
  geom_vline(
    aes(xintercept = first_collection_date_dug),
    linetype = "dotted",
    color = "#c44601",
    linewidth = 2
  ) +
  
  facet_wrap(~year, scales = "free_x") +
  
  theme_bw() +
  theme(
    strip.text = element_text(face = "bold"),
    axis.text.x = element_text(face = "bold", angle = 45, hjust = 1, , size = 10),
    axis.text.y = element_text(face = "bold", size = 10),
    panel.grid.major = element_blank(),
    panel.grid.minor = element_blank()
  ) +
  
  labs(
    x = "Date",
    y = "Cumulative Degree Days"
  )
```

Richardson County

```
ggplot(dd_plot_data_ric, aes(x = date, y = CDD)) +
  geom_line(color = "black", linewidth = 0.8) +
  
  # DD threshold line
  geom_vline(
    aes(xintercept = dd_threshold_date_ric),
    linetype = "dashed",
    color = "#029356",
    linewidth = 0.8
  ) +
  
  # First emergence line
  geom_vline(
    aes(xintercept = first_emergence_date_ric),
    linetype = "solid",
    color = "#2546f0",
    linewidth = 0.8
  ) + 
  # First sampling
  geom_vline(
    aes(xintercept = first_collection_date_ric),
    linetype = "dotted",
    color = "#c44601",
    linewidth = 2
  ) +
  
  facet_wrap(~year, scales = "free_x") +
  
  theme_bw() +
  theme(
    strip.text = element_text(face = "bold"),
    axis.text.x = element_text(face = "bold", angle = 45, hjust = 1, , size = 10),
    axis.text.y = element_text(face = "bold", size = 10),
    panel.grid.major = element_blank(),
    panel.grid.minor = element_blank()
  ) +
  
  labs(
    x = "Date",
    y = "Cumulative Degree Days"
  )
```
